# Using machine learning to dissect host kinases required for *Leishmania* internalization and development

**DOI:** 10.1101/2024.05.16.593986

**Authors:** Ling Wei, Umaru Barrie, Gina M. Aloisio, Francis T. H. Khuong, Nadia Arang, Arani Datta, Alexis Kaushansky, Dawn M. Wetzel

## Abstract

The *Leishmania* life cycle alternates between promastigotes, found in the sandfly, and amastigotes, found in mammals. When an infected sandfly bites a host, promastigotes are engulfed by phagocytes (*i.e.*, neutrophils, dendritic cells, and macrophages) to establish infection. When these phagocytes die or break down, amastigotes must be re-internalized to survive within the acidic phagolysosome and establish disease. To define host kinase regulators of *Leishmania* promastigote and amastigote uptake and survival within macrophages, we performed an image-based kinase regression screen using a panel of 38 kinase inhibitors with unique yet overlapping kinase targets. We also targeted inert beads to complement receptor 3 (CR3) or Fcγ receptors (FcR) as controls by coating them with complement/C3bi or IgG respectively. Through this approach, we identified several putative host kinases that regulate receptor-mediated phagocytosis and/or the uptake of *L. amazonensis*. Findings included kinases previously implicated in *Leishmania* uptake (such as Src family kinases (SFK), Abl family kinases (ABL1/c-Abl, ABL2/Arg), and spleen tyrosine kinase (SYK)), but we also uncovered many novel kinases. Our methods also predicted host kinases necessary for promastigotes to convert to amastigotes or for amastigotes to survive within macrophages. Overall, our results suggest that the concerted action of multiple interconnected networks of host kinases are needed over the course of *Leishmania* infection, and that the kinases required for the parasite’s life cycle may differ substantially depending on which receptors are bound and the life cycle stage that is internalized. In addition, using our screen, we identified kinases that appear to preferentially regulate the uptake of parasites over beads, indicating that the methods required for *Leishmania* to be internalized by macrophages may differ significantly from generalized phagocytic mechanisms. Our findings are intended to be used as a hypothesis generation resource for the broader scientific community studying the roles of kinases in host-pathogen interactions.

## 1. Introduction

*Leishmania* is an obligate intracellular parasite of phagocytic cells that causes either cutaneous or visceral disease, depending on the infecting parasite species and the immunological status of the host [1, 2]. Since millions of patients are affected worldwide, and drugs for leishmaniasis are both ineffective and have notable host toxicity, new anti-leishmanial drugs are needed. Further research into the mechanisms driving *Leishmania* pathogenesis is essential to identify new avenues for therapeutic development [3]. Infection of phagocytes is initiated by *Leishmania* promastigotes, the form that develops in the female sandfly’s digestive tract. Infection of macrophages also can occur if the amastigotes that replicate within phagolysosomes of mammalian host cells are found outside phagocytes (*i.e.*, if the mammalian cells die or break down) [4]. Several host cell receptors on macrophages have previously been identified as critical in *Leishmania* binding, internalization, and subsequent survival within host cells [5]. As described in the literature, the fibronectin receptor (integrin α5β1) [6], mannose receptor [7], third complement receptor CR3 [8], and first complement receptor CR1 [9] facilitate promastigote identification. Additionally, CR3, heparin binding protein, and the Fc receptor (FcγR) mediate amastigote uptake by phagocytes [10]. Ligation of these specific receptors is known to elicit downstream activation of signaling pathways necessary for actin polymerization and parasite internalization.

Overall, there has been a prevailing belief in the *Leishmania* research community that early signaling events involved in *Leishmania* internalization closely resemble other phagocytic mechanisms [7, 22]. This belief has largely stemmed from older studies that focused on host cell responses to cytoskeleton-active agents [7]. This hypothesis contrasts with other intracellular parasites such as *Toxoplasma* and *Trypanosoma cruzi*, where cell invasion is predominantly reliant on parasitic machinery [3]. However, even if this is the case, binding of different receptors may trigger distinct downstream host signaling pathways that regulate *Leishmania* survival and intracellular development, based on previously published data. For example, *L. amazonensis* binding to CR3 is known to trigger Rho GTPase recruitment and cytoskeletal rearrangement. Phagocytic vesicles formed post-CR3-mediated uptake exhibit a snug fit, and phagolysosome maturation is delayed, which accompanies impaired accumulation of LAMP-1 and cathepsin-D [8]. CR3 binding also hampers IFN-γ-induced pro-inflammatory signaling and reduces IL-12-mediated H_2_O_2_ production [8]. Conversely, when *L. amazonensis* binds FcR, it triggers Rac GTPase recruitment and cytoskeletal rearrangement. Rac, in turn, activates NADPH oxidase, and the resulting phagocytic vesicles formed after FcR-mediated uptake are expansive [11]. Otherwise, many of the mechanisms by which *Leishmania* modulates signaling pathways and their contribution to the phagocytosis process have yet to be elucidated [3].

Our ongoing efforts have been directed at identifying the host cell signaling pathways that permit *Leishmania* internalization by mammalian macrophages. Furthermore, we have hypothesized that targeting the host-based process through which both life cycle stages of *Leishmania* are internalized by phagocytic cells represents a potential therapeutic strategy, and that there are aspects of *Leishmania* uptake that differ from general phagocytic mechanisms. By blocking the uptake process, the initial infection step by promastigotes and subsequent spread of *Leishmania* by amastigotes within the host could be prevented [6–10, 12, 13]. To this end, we have employed a multi-color immunofluorescence assay to define *L. amazonensis* promastigotes or amastigotes that have been internalized by, or are surviving within, macrophages [12–14]. By directing our experiments at known signaling partners of previously-identified kinases in other biological systems, we have defined a Src-Abl/Arg-Syk related signaling pathway that governs *Leishmania* uptake and resulting disease manifestations in mouse models [13, 15–17]. Such studies have been fruitful but are limited by what is known about interactions with each identified kinase in other systems. Employing a broader yet systematic screening approach would provide a more complete picture of the signaling pathways used by *Leishmania* to enter and survive in macrophages.

We have previously used a machine learning approach named kinase regression (KiR) to identify host kinases responsible for malaria liver-stage infection [18], blood-brain barrier integrity [19, 20] and dengue infection in hepatocytes [21]. By incorporating cellular phenotypic data from a small panel of kinase inhibitors with existing biochemical data of these inhibitors against 291 human protein kinases, KiR can make predictions on the key kinase regulators driving that phenotype using an elastic net regularization algorithm [22].

Therefore, in this study, we used KiR to identify the most important host kinase regulators of *L. amazonensis* uptake and development. A small subset of these predictions was empirically confirmed. Collectively, our predictions have identified important host cell regulators of *L. amazonensis* infection, which vary depending on the entry receptor and form of the parasite. These findings are shared here for the broader parasitology community and offer promising insights into defining effective host-based targets for antiparasitic treatment.

## 2. Materials and methods

### 2.1 Mice

C57BL/6 mice were purchased from Jackson Laboratory (Bar Harbor, ME). The Institutional Animal Care and Use Committee at UT Southwestern approved all experimental protocols.

### 2.2 Cell culture

For the initial kinase screen, cells were harvested from the tibias and femurs of mice and differentiated into bone-marrow-derived (BM) primary macrophages over 7 days by growing them in DMEM supplemented with 10% FBS and 20% supernatant from L929 cells.

Differentiation into BM macrophages was confirmed previously by fluorescence-activated cell sorter (FACS) using antibodies to F4/80 and C11b (eBioscience, San Diego, CA); over 98% of differentiated cells were positive for both markers. For confirmation screens with trametinib, RAW 264.7 cells (American Type Culture Collection), a murine macrophage-like cell line, was used.

### 2.3 Parasite culture

*Leishmania amazonensis* promastigotes (strain MHOM/BR/767/LTB0016) were grown at 26°C in Schneider’s Drosophila medium supplemented with 15% heat-inactivated, endotoxin-free FBS and 10 μg/ml gentamicin. For macrophage invasion, promastigote cultures were incubated at stationary phase for 5-7 days to maximize infective metacyclic promastigotes and defined as such using a Percoll gradient [12]. *L. amazonensis* amastigotes were grown axenically (outside of macrophages) according to [23, 24]; amastigotes were grown at 32°C in M199 (Invitrogen) at pH 4.5 supplemented with 20% FBS, 1% penicillin-streptomycin, 0.1% hemin (25 mg/ml in 50% triethanolamine), 10 mM adenine, 5 mM L-glutamine, 0.25% glucose, 0.5% Trypticase Soy Broth [Sigma], and 40 mM sodium succinate.

### 2.4 Phagocytosis assays

Primary macrophages were incubated overnight in M-CSF-starved media. One hour prior to phagocytosis, macrophages were activated with phorbol myristate acetate (PMA, Sigma) for 1 h. Experiments were performed at ∼50% confluence. For drug screening experiments, macrophages were preincubated in 500 nM of each kinase inhibitor or dimethyl sulfoxide (DMSO, Sigma) for 2 h. For experiments with C3bi-opsonized particles, 2-μm-diameter latex yellow-green beads (Sigma) were coated with human IgM (Sigma) at 1:100 for 1 h at 37°C. They were then incubated in freshly isolated mouse serum diluted 1:2 in phosphate-buffered saline (PBS) for 1 h at 37°C, which allowed C3bi opsonization. For IgG experiments, yellow-green beads were incubated with rabbit IgG (Sigma) diluted 1:250 for 1 h at 37°C. Macrophages were incubated with a ratio of 10 to 15 particles per cell for the indicated timepoints at 37°C in the presence of inhibitors. For survival assays of 24 and 72 h, macrophages were washed with fresh media to remove any beads that had not been taken up and then given fresh media with inhibitors. As experiments were performed in 96 well plates, 25% formaldehyde was added directly to wells to make a final dilution of 3% formaldehyde. Samples were fixed with 3% formaldehyde for 15 min, and blocked with 2% bovine serum albumin (BSA) without permeabilization. Samples with C3bi-coated beads were incubated with rabbit anti-human IgM (Sigma) at 1:250. All samples were labeled with A568-conjugated goat anti-rabbit secondary antibody (Invitrogen) at 1:250. Samples were permeabilized with 0.25% Triton-X100 (Sigma) for 10 min and rinsed with PBS. Samples were rinsed with PBS again and finally stained with 4′,6-diamidino-2-phenylindole (DAPI, Invitrogen). This process, termed a “three-color immunofluorescence assay”, allowed us to distinguish the beads that were internalized from those that remained outside cells. A full protocol for this process can be found in [14].

### 2.5 *Leishmania* uptake assays

Primary mouse bone marrow-derived macrophages were employed for all assays. Macrophages were grown on black 96-well plates and were incubated with 500 nM kinase inhibitors (screening assays) or 1 μM trametinib (confirmation assays) for 2 h, after which macrophages were washed with media followed by parasite addition to macrophages/media/inhibitors. Macrophages were also activated with PMA for 1 h prior to experiments. For C3bi opsonization experiments, promastigotes and amastigotes were incubated with freshly isolated mouse serum for 1 h. For IgG opsonization, amastigotes were coated with anti-P8–proteoglycolipid complex (monoclonal antibody IgG1) for 1 h. Because promastigotes are unlikely to encounter specific antibodies during an animal infection (it is not a biologically relevant condition if an animal has not been previously infected), IgG opsonization of promastigotes was not performed. Promastigotes were incubated at a ratio of 10 parasites: 1 macrophage and amastigotes were incubated at a ratio of 5 parasites: 1 macrophage. For experiments performed in 96-well plates, 25% formaldehyde was added directly to wells to make a final dilution of 3% formaldehyde. External promastigotes were labeled with mouse anti-gp46, while external amastigotes were labeled with mouse anti-P8. All coverslips and plates were labeled with A594-conjugated goat anti-mouse secondary antibody (Invitrogen). After permeabilization, promastigotes were relabeled with mouse anti-gp46, and amastigotes were relabeled with mouse anti-P8. Either life cycle stage was also labeled with A488-conjugated goat anti-mouse secondary antibody (Invitrogen); samples were co-stained with DAPI. For all experiments, at least 10 randomly selected, distinct fields were visualized containing at least 100 macrophages and at least 100 parasites per experimental determination. All experiments were performed 3 times unless otherwise indicated.

### 2.6 Parasite survival assays

For parasite survival assays, macrophages were treated as above, except longer timepoints of 24 h and 72 h were employed. At the 24 h mark, cells were washed with media, and fresh media (with inhibitors) was given. For experiments on coverslips, macrophages were fixed with 3% formaldehyde and for experiments performed in 96-well plates, 25% formaldehyde was added directly to wells to make a final dilution of 3% formaldehyde. After permeabilization, promastigotes had switched to amastigotes based on morphological changes and binding of amastigote-specific antibodies, and amastigotes were labeled with mouse anti-P8 and A488-conjugated goat anti-mouse secondary antibody (Invitrogen); samples were co-stained with DAPI. For all experiments, at least 10 randomly selected, distinct fields were visualized containing at least 100 macrophages and at least 100 parasites per experimental determination. All experiments were performed 3 times unless otherwise indicated.

### 2.7 Microscopy

For the initial kinase screen, fluorescence was visualized in the UT Southwestern High Throughput Screening Core on a InCell Analyzer 6000 (GE Healthcare Life Sciences) at 20X. At least 10 randomly selected fields from multiple technical replicates were visualized for a total of at least 100 macrophages per experimental determination. The mean phagocytic index (PI), the number of particles internalized per 100 mammalian cells, was calculated for each experiment using a pipeline from CellProfiler (https://cellprofiler.org, Broad Institute). The phagocytic index for control cells was taken as the maximum value (100%) for each experiment. Means ± standard errors (SE) were calculated, and each biological experiment was performed at least three times. Confirmation assays using trametinib were performed similarly, except that samples were visualized on a Cytation 5 (BioTek), and analysis was performed using the system’s automated software [12]. One-sample Student’s *t*-tests were performed to assess statistical significance.

### 2.8 Kinase regression

The elastic net regularization algorithm used for this study was published previously [19]. The kinase regression (KiR) approach exploits the polypharmacology of a small panel of kinase inhibitors and makes predictions on kinases important for specific cellular phenotypes and the effect of untested kinase inhibitors on the phenotypes based on the linear combination of the contributions of kinases to the phenotypes [22]. For this study, the mean number of internal parasites per 100 mammalian cells upon kinase inhibitor treatment was normalized to the phagocytic index upon DMSO treatment as control. The normalized phagocytic indices, together with the biochemical data of the panel of 38 kinase inhibitors against 291 recombinant protein kinases, were input into the elastic net regularization algorithm with a condition-specific cross-validation strategy. The glmnet package (https://github.com/bbalasub1/glmnet_python, version 2.2.1) in Python (https://www.python.org, version 3.7.6) was used to perform elastic net regularization, with the elastic net mixing parameter of 0.8.

### 2.9 Network generation

To build the phosphosignaling network for a set of KiR-predicted kinases, the shortest paths between any pair of kinases within that set was identified using the kinase-substrate phosphorylation database on PhosphoSitePlus^®^ (https://www.phosphosite.org/staticDownloads) as the background network, and all the paths were combined to form the phosphosignaling network that describes the paths through which the signals may propagate. Searches for the shortest paths between kinases were done using NetworkX (https://github.com/networkx/networkx), a Python package for analyzing complex network structures, such as analyzing structure and dynamics of networks, generating networks of various types, building network models, and designing new network algorithms. The shortest paths were computed by function “*shortest_path*” and the shortest path lengths were computed by function “*shortest_path_length*” in NetworkX.

### 2.10 Visualization

Schematics were created with BioRender.com. Phosphosignaling networks were visualized using Cytoscape (https://cytoscape.org, version 3.9.1), with the hierarchic layout from yFiles layout algorithms (https://www.yworks.com/products/yfiles-layout-algorithms-for-cytoscape).

## 3. Results

### 3.1 A broad-scale, overlapping-target kinase inhibitor screen implicates many host cell kinases in *Leishmania* uptake of macrophages and intracellular development

As indicated above, multiple surface protein receptors, including complement receptor CR3, facilitate promastigote uptake; both CR3 and Fc receptor (FcR) permit amastigote uptake [25–27]. We hypothesized that if *Leishmania* promastigotes and amastigotes bound different extracellular receptors prior to internalization, these receptors might influence the signaling pathways facilitating parasite uptake by and survival within macrophages. To test this hypothesis and study the effects of binding different receptors, we cultured *L. amazonensis* promastigotes and axenic amastigotes [13, 25, 26]; promastigotes were opsonized with C3bi and amastigotes were coated with C3bi or IgG (Fig. 1A). We did not coat promastigotes with IgG because antibody opsonization of promastigotes is unlikely to be physiologically relevant (unless a host organism were infected with *Leishmania* parasites multiple times). We also coated yellow-green beads with C3bi or IgG as controls. We simultaneously pre-treated bone marrow-derived macrophages with a panel of 38 kinase inhibitors that target multiple different kinases throughout the mammalian kinome; inhibitors remained in place throughout the experiment. Cognate kinase targets inhibited by these drugs and their biochemical data against 291 recombinant kinases can be found in Table S1. Four different co-incubation timepoints were employed to broadly characterize *Leishmania*/bead uptake into and survival within macrophages: 0.5 hour for initial parasite uptake, 3 hours to encapsulate the end of parasite uptake and trafficking to the parasitophorous vacuole (PV), 24 hours to demonstrate transition from promastigote to amastigote forms, and 72 hours to assess intracellular replication of amastigotes (Fig. 1B) before fixation with formaldehyde. We then processed samples for immunofluorescence using a published method that allows us to differentiate extracellular and intracellular parasites. Briefly, we first labeled external parasites red prior to sample permeabilization, and then labeled all parasites green after permeabilization (Fig. 1C) [14]. We found that our 38 screened kinase inhibitors had a diverse impact on *Leishmania* uptake and development in macrophages at different timepoints, life cycle stages, and when parasites had bound different cell surface receptors (Fig. 1D, Fig. S1A and B, Table S2). Use of this set of kinase inhibitors confirmed prior data from our laboratory and others indicating that the Abl/Arg inhibitors imatinib and dasatinib, and the combination Src/Abl/Arg inhibitor bosutinib limited *Leishmania* and bead uptake at 30 minutes [12, 13, 15, 28]. Similarly, uptake of beads was decreased by published phagocytic inhibitors, such as our ROCK inhibitor (Y-27632) [29]. However, a number of other kinase inhibitors that we tested also affected internalization or survival of *Leishmania* promastigotes and amastigotes (Fig. 1D, Table S2), as well as the uptake of coated beads (Fig S1A).

**Fig. 1.**
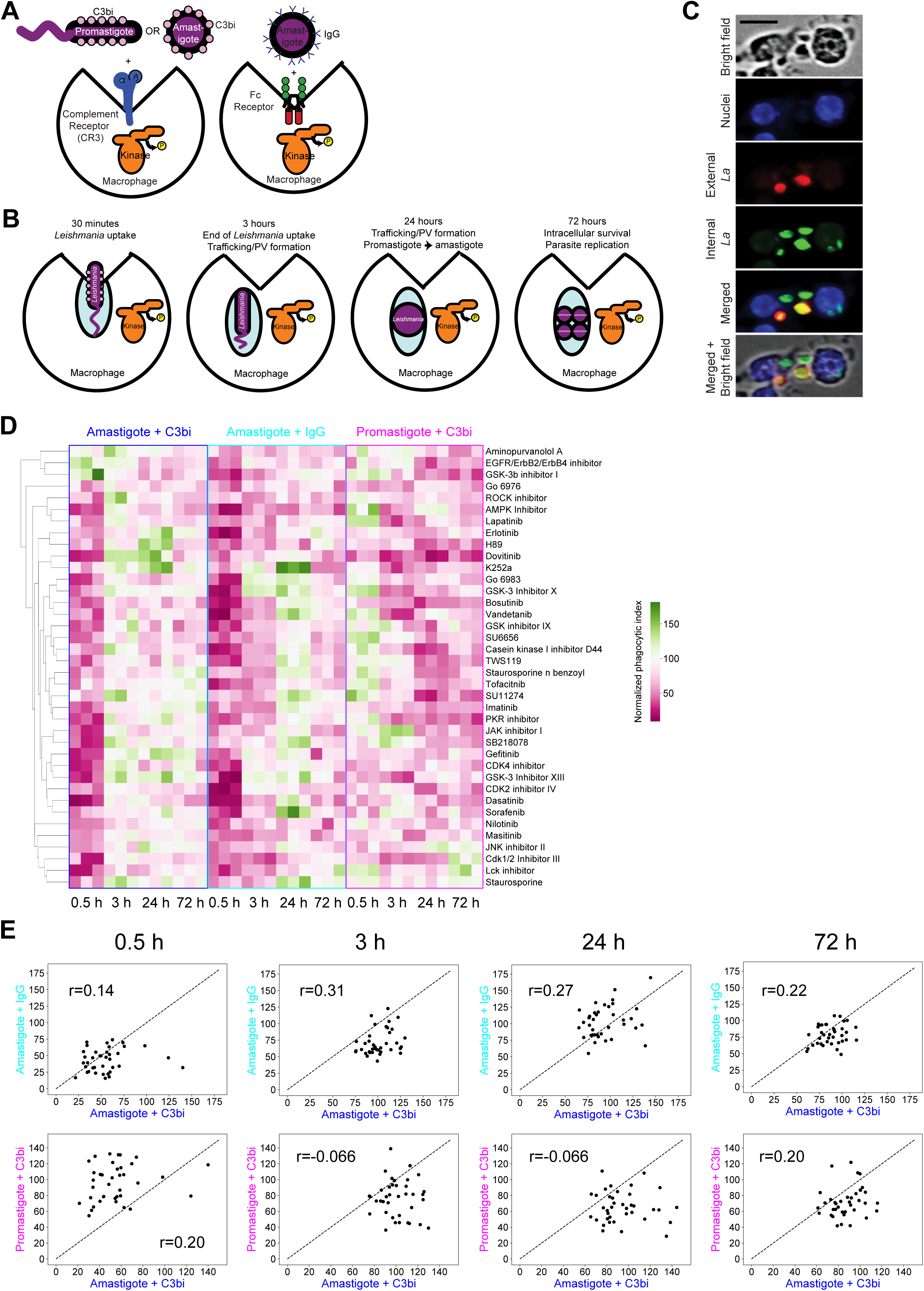
Host-targeted kinase inhibitors modulate infection of *Leishmania* in macrophages. **(A)** Invasion of *Leishmania* parasites is initiated by C3bi-opsonized promastigotes or amastigotes interacting with complement receptors (*e.g*., CR3) on macrophages, or amastigotes entering macrophages via an IgG-mediated mechanism by binding to FcR. **(B)** Four representative stages of *Leishmania* infection in macrophages were studied: 30 min to characterize initial parasite uptake, 3 h to assess the end of parasite uptake and trafficking to the parasitophorous vacuole (PV), 24 h to visualize the transition from promastigote form to the amastigote form, and 72 h to identify kinases needed for intracellular replication of amastigotes. **(C)** Microscopy images showing that our multicolored fluorescence assay allows labeling of all parasites (green channel) and parasites outside of macrophages (red channel). DAPI was used to label cell nuclei (blue channel). See Materials and Methods and [14] for details on fluorescent labeling and microscopy. **(D)** Cluster map showing the impact of a panel of 38 kinase inhibitors on *Leishmania* infection in macrophages. *Leishmania* infection was quantified by phagocytic index - the number of internalized particles per 100 mammalian cells, at 0.5, 3, 24, and 72 h post infection. Phagocytic indices for kinase inhibitor-treated cells were normalized to the phagocytic index of DMSO-treated control cells. Three independent experiments were performed. **(E)** Correlations of normalized phagocytic index (average of all three biological replicates) upon kinase inhibitor treatment between amastigote-initiated infection via C3bi- and IgG-mediated mechanisms (top) and between amastigote- and promastigote-initiated infection via C3bi-mediated mechanism (bottom). Pearson correlation coefficient (r) is shown in each plot.

### 3.2 Based on initial screening results*, Leishmania* uptake and intracellular development appears to be differentially impacted by host kinases depending upon life cycle stage and the entry receptor

To examine how inhibition of various kinases varied outcomes under different entry conditions, we assessed correlations of phagocytic indexes and the numbers of parasites surviving within macrophages over time across the panel of 38 kinase inhibitors between amastigote uptake via CR3 and FcR, and between promastigote and amastigote uptake via CR3. Interestingly, for these 38 kinase inhibitors, we observed little association (r: 0.14 – 0.31) between the inhibition of C3bi- and IgG-mediated entry mechanisms over time (Fig. 1E, top), suggesting that binding of different receptors triggered different uptake and survival mechanisms. We also found little correlation (r: -0.066 – 0.20) between promastigote and amastigote forms entering macrophages via a C3bi-mediated mechanism (Fig. 1E, bottom), suggesting that receptor binding was not the only thing that stimulated differential pathways regulating parasite uptake and survival.

### 3.3 Kinase regression predicts that differential kinase regulatory mechanisms are required for *Leishmania* infection of macrophages compared to phagocytosis of inert beads

Overall, there has been a prevailing belief in the *Leishmania* research community that early signaling events involved in *Leishmania* internalization closely resemble generalized phagocytic mechanisms [7, 22], although this historical belief has been evolving over time. Instead, given the unique pattern of kinase inhibitor effects observed across different entry mechanisms and parasite forms (Fig. 1D, E), we hypothesized that distinct kinase regulatory processes would regulate C3bi or IgG-opsonized bead uptake (generalized phagocytosis) compared to promastigote or amastigote infection (*Leishmania* internalization). Therefore, next, we sought to understand the similarities and differences between the kinase regulatory program governing host control of *Leishmania* infection and the mechanisms that regulate phagocytosis in general.

To do so, we used a machine learning approach called kinase regression (KiR), which utilizes the kinase screen data on *Leishmania* infection along with existing biochemical data of these kinase inhibitors against 291 human kinases (Table S1, Fig. S2) [30] to make predictions on the kinases important for *Leishmania* infection of macrophages (Fig. 2A). For each experimental condition, KiR was performed on the data taken at the respective stages of infection. Known and novel kinase regulators were predicted to modulate *Leishmania* invasion and intracellular development. A full list of kinases predicted to facilitate or inhibit bead and *L. amazonensis* uptake and persistence is shown in Table S3.

**Fig. 2.**
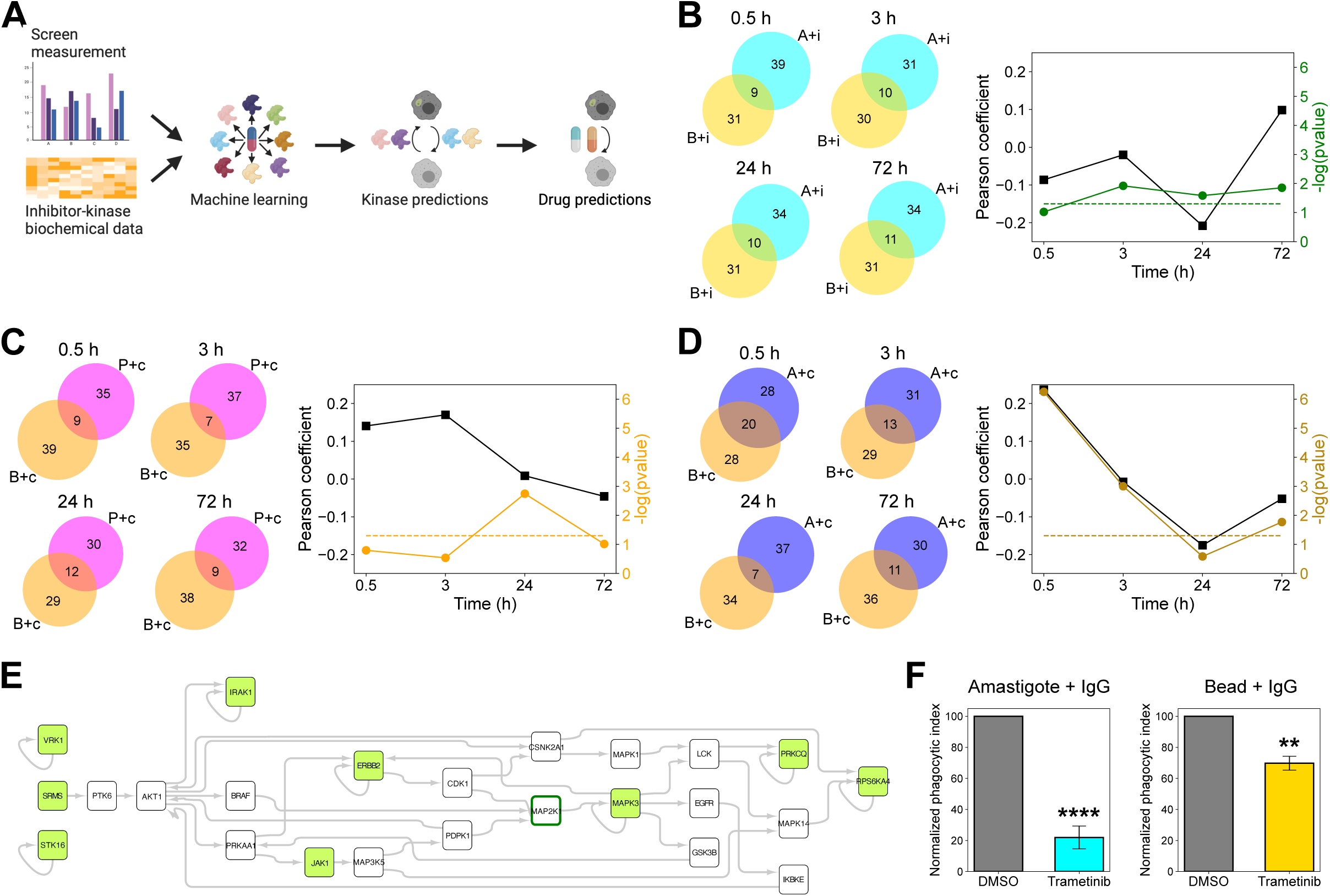
Host kinases regulate *Leishmania* infection through mechanisms independent of phagocytosis. **(A)** Workflow of using our kinase inhibitor screen and a machine learning algorithm to identify host kinase regulators important for *Leishmania* infection in macrophages. **(B)** Left: Venn diagrams showing the number of kinases predicted to modulate IgG-mediated uptake of amastigotes and beads at different stages of infection. Right: black squares represent the Pearson correlation coefficients calculated from the normalized phagocytic index upon kinase inhibitor treatment between amastigote-initiated infection via C3bi- and IgG-mediated mechanisms; green circles represent the p-values from hypergeometric test performed on KiR-predicted kinases between amastigote and bead uptake via IgG-mediated mechanism; green dashed line corresponds to p-value of 0.05. **(C)** Left: Venn diagrams showing the number of kinases predicted to modulate C3bi-mediated uptake of promastigotes and beads at different stages of infection. Right: black squares represent the Pearson correlation coefficients calculated from the normalized phagocytic index upon kinase inhibitor treatment between promastigote-initiated infection and bead uptake via a C3bi-mediated mechanism. Orange circles represent the p-values from hypergeometric test performed on KiR-predicted kinases between promastigote and bead uptake via C3bi-mediated mechanism. Orange dashed line corresponds to p-value of 0.05. **(D)** Left: Venn diagrams showing the number of kinases predicted to modulate C3bi-mediated uptake of amastigotes and beads at different stages of infection. Right: black squares represent the Pearson correlation coefficients calculated from the normalized phagocytic index upon kinase inhibitor treatment between amastigote-initiated infection and bead uptake via C3bi-mediated mechanism. Brown circles represent the p-values from hypergeometric test performed on KiR-predicted kinases between amastigote and bead uptake via C3bi-mediated mechanism. Brown dashed line corresponds to p-value of 0.05. **(E)** Phosphosignaling network built from the overlapping kinases predicted in IgG-mediated uptake of amastigotes and beads at 3-hour time point. Kinases predicted by KiR are in green. **(F)** Targeting the ERK pathway using a MEK inhibitor (trametinib) reduced macrophage uptake of amastigotes via an IgG-mediated mechanism (left), as well as the phagocytosis of IgG-coated beads (right). The phagocytic index, the number of intracellular amastigotes or beads per 100 macrophages, was calculated for each trametinib-treated biological replicate and normalized to the DMSO-treated biological replicate (100%). Shown is the normalized phagocytic index at 3-hour post internalization (n = 4 biological replicates for amastigotes; 3 biological replicates for beads). One of trametinib’s targets, MAP2K1/MEK1, is highlighted in dark green in (E).

To study how signaling events change over time for this intracellular parasite versus coated beads, we compared KiR predictions between *Leishmania* and bead uptake by, and persistence within, macrophages. We reasoned that three distinct options were feasible. (1) If early infection was mediated strictly by host regulators of phagocytosis, we would observe a high degree of overlap in kinases predicted to regulate C3bi-coated bead versus parasite uptake and IgG-coated bead versus amastigote uptake, which would then diverge as parasites changed their host requirements at later (survival) time points. (2) In contrast, if significant overlap existed throughout infection at both early and late timepoints, we reasoned that phagocytic uptake would intrinsically shape the host regulators of infection throughout the process. (3) Finally, if little overlap in host kinases for parasites versus beads were present early on during infection, we reasoned that distinct mechanisms would dominate entry.

Interestingly, we observed examples of all three patterns described above. Kinase predictions of IgG-mediated amastigote and bead uptake suggested modest overlap through the entire parasite lifecycle at early and late time points (Fig. 2B), most similar to model (2) above. Despite this, most KiR-predicted kinases were unique to amastigote + IgG infection throughout the time course. However, when comparing promastigotes and beads treated with C3bi, we observed less overlap between the kinase predictions than would be predicted by chance (Fig. 2C). We interpret this finding to suggest that C3bi-promastigotes have largely unique kinase regulatory requirements from beads that enter strictly by phagocytosis, more similar to the model described in point (3) above. Finally, and perhaps most strikingly, we saw a dramatic similarity in the overlap between kinases predicted to regulate early time points of amastigotes and beads treated with C3bi. This overlap waned as time progressed, most similar to model (1) above (Fig. 2D). The significant overlap of kinase regulators between C3bi-amastigote and bead uptake suggested that internalization of amastigotes via a C3bi-mediated mechanism may be more phagocytosis-driven than other methods. However, once an amastigote successfully invades macrophages, non-phagocytic-associated signaling may become essential for its intracellular development. These results suggested that the mechanism and receptor driving the initial uptake of *Leishmania* parasites often but does not always influence its ultimate cellular fate, which is consistent with a subset of prior literature [9, 11].

### 3.4 A subset of kinases and kinase signaling pathways are involved in both *Leishmania* infection of macrophages and phagocytosis

To better understand the portion of network-level kinase signaling that overlapped with phagocytosis during *Leishmania* infection, we built phosphosignaling networks from the predicted kinases overlapping between *Leishmania* and bead uptake conditions, using the kinase-substrate phosphorylation database on PhosphoSitePlus^®^ as the background network (Fig. 2E). PhosphoSitePlus^®^ is a dynamically updated knowledgebase that integrates both low- and high-throughput data sources into a single reliable and comprehensive resource for studying experimentally observed post-translational modifications in the regulation of biological processes [31]. The kinase phosphosignaling networks that we built revealed many additional kinases that were not predicted by KiR but were predicted to serve as intermediates, connecting KiR-predicted kinases. For instance, at 3 h of infection where amastigotes have finished internalizing into the cell and start trafficking to PV, our model predicted that MAPK signaling plays an important role in amastigote-initiated uptake as well as IgG-bead uptake (Fig. 2E). Furthermore, a canonical MAPK signaling architecture (MAP3K-MAP2K-MAPK) was also implicated in the network built from the kinases shared between promastigote and bead uptake via a C3bi-mediated mechanism at 3 h of infection (Fig. S3A). Similarly, another canonical MAPK signaling architecture, MAP2K-MAPK-MAPKAPK, was inferred by the network built from the shared kinases between promastigote and bead uptake at the early stage of invasion (Fig. S3B). To experimentally evaluate these predictions, we assessed infection of macrophages by IgG-opsonized amastigotes and IgG-coated beads at multiple timepoints in the presence of trametinib, a MEK (MAP2K) inhibitor (3 h data shown in Fig. 2F; additional timepoints and data for promastigotes can be found in [17]). We found that inhibiting MEK decreased macrophage infection by *Leishmania* amastigotes, as well as the uptake of beads. Overall, our network analysis suggests that a subset of phagocytic pathways is required for *Leishmania* uptake and survival. However, there are more non-phagocytic-associated signaling pathways essential for establishing *Leishmania* infection in macrophages, which are analyzed further in the following sections.

### 3.5 Both differential and overlapping kinase signaling pathways are predicted to be associated with amastigote versus promastigote infection of macrophages

Promastigotes and amastigotes differ in their morphology, proteomes, and membrane constituents. Given the distinct origins and characteristics of these two parasite life cycle stages, it would be reasonable to hypothesize that different sets of host kinase regulators are involved in the uptake and subsequent regulation of parasite survival within the host cell. Our KiR model indeed predicted the involvement of different kinases for promastigote entry into macrophages compared to amastigotes, including several novel kinases not previously implicated in *Leishmania* infection and kinases unrelated to phagocytosis (Fig. 3A). Interestingly, a small set of kinases were predicted to be important for intracellular development for both promastigotes and amastigotes via a C3bi-mediated entry mechanism, and this overlap was statistically significant compared to random selections from the pool of 291 kinases in the screen (Fig. 3A). Since promastigotes transition to amastigotes approximately 24 h after macrophage infection, the increased overlap of predicted kinases between promastigote- and amastigote-initiated infection at 24 h and 72 h post-infection may be attributed to this transition within macrophages at later infection stages. Nonetheless, the significant number of kinases predicted to be uniquely essential for promastigote- or amastigote-initiated infection suggests that distinct kinase signaling pathways regulate the establishment of *Leishmania* parasites in macrophages during and after life cycle stage conversion.

**Fig. 3.**
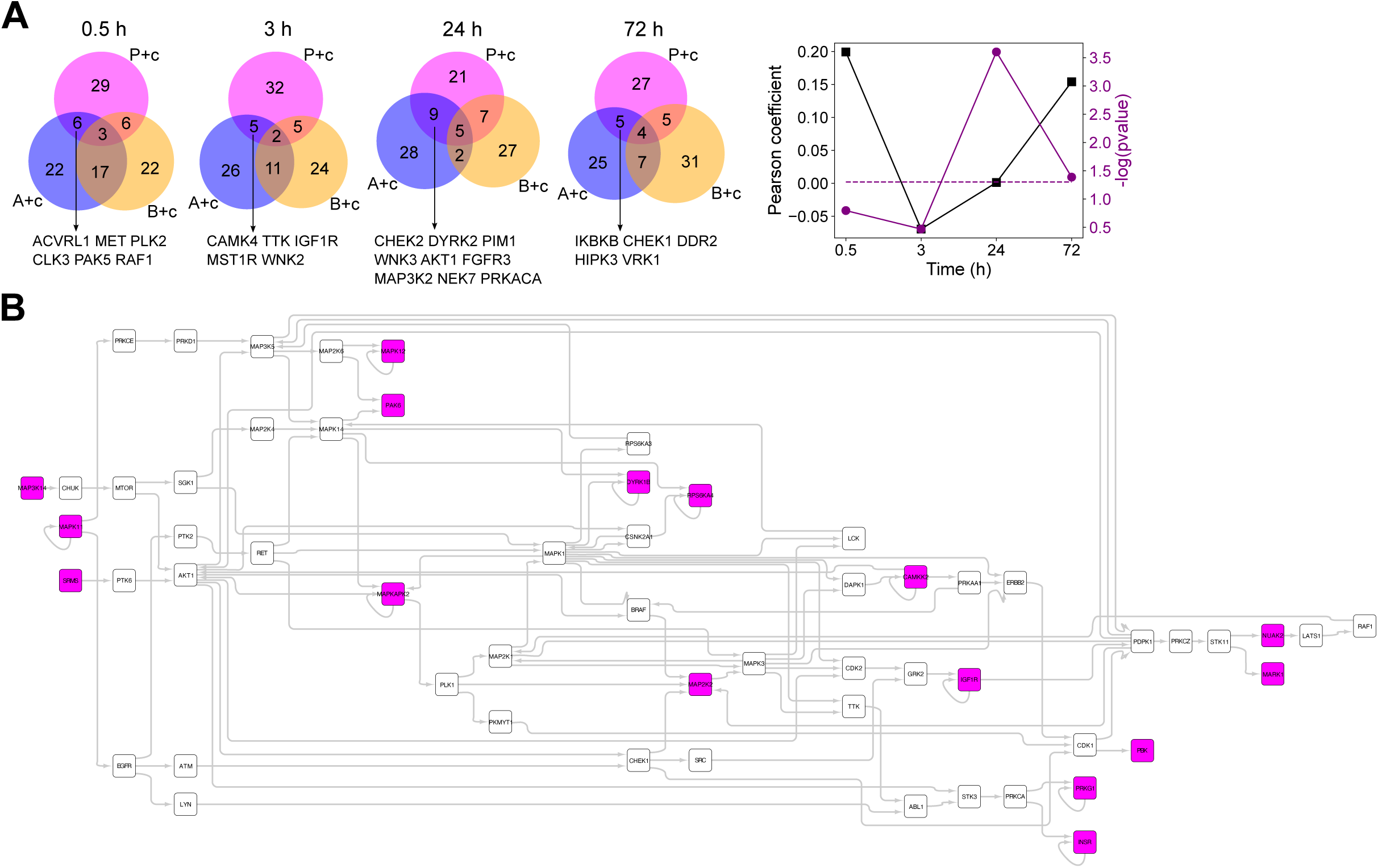
Host kinases differentially regulate *Leishmania* promastigote and amastigote infection in macrophages. **(A)** Left: Venn diagrams showing the number of kinases predicted to modulate C3bi-mediated uptake of promastigotes, amastigotes, and beads at different stages of infection. Kinases predicted to regulate both promastigote and amastigote infection but not bead uptake are listed. Right: black squares represent the Pearson correlation coefficients calculated from the normalized phagocytic index upon kinase inhibitor treatment between promastigote- and amastigote-initiated infection via C3bi-mediated mechanism; purple circles represent the p-values from hypergeometric test performed on KiR-predicted kinases between promastigote and amastigote infection via C3bi-mediated mechanism; purple dashed line corresponds to p-value of 0.05. **(B)** Phosphosignaling network built from the kinases predicted to exclusively regulate promastigote-initiated infection at 24 h post infection (labeled in magenta), using the kinase-substrate phosphorylation database on PhosphoSitePlus^®^ as the background network.

To further capture the molecular details of kinase signaling that are unique to promastigote-initiated infection, we built phosphosignaling networks from KiR-predicted kinases unique to the uptake of promastigotes but not uptake of amastigotes or beads via a C3bi-mediated entry mechanism. For example, at 24 h post infection, when internalized promastigotes transition to amastigotes, MAPK signaling was once again enriched in a network built from kinases predicted to exclusively regulate promastigote-initiated infection, with 12 out of 58 kinases in this network belonging to the MAP kinase family (Fig. 3B, hypergeometric test p-value of 0.0046). This pathway analysis suggests that different kinase signaling pathways are required for establishing promastigote- and amastigote-initiated infection in macrophages, which is consistent with prior literature characterizing differences between these signals [9].

### 3.6 Differential kinase signaling is predicted to drive C3bi-versus IgG-mediated amastigote uptake and survival

We next inquired if differential kinase signaling would mediate C3bi-versus IgG-mediated amastigote uptake and survival. Comparing the KiR-predicted kinase regulators between C3bi- and IgG-amastigotes during infection, we observed limited overlap in the regulatory kinases predicted in each condition (Fig. 4A). In fact, overlap between predicted regulatory kinases at 3 and 24 h post-infection was less than would be predicted by chance alone. These KiR predictions support the hypothesis that the uptake of amastigotes through a C3bi-mediated mechanism engages a different set of host kinases compared to those regulating infection through an IgG-mediated mechanism. We then utilized our data to identify putative regulatory C3bi-versus IgG-mediated factors essential exclusively for *L. amazonensis* infection. To achieve this, we compared our KiR-predicted kinase regulators between C3bi- and IgG-mediated uptake of inert beads as controls (Fig. 4B). As anticipated, akin to the uptake of amastigotes, distinct sets of kinases were implicated in regulating C3bi- and IgG-mediated phagocytosis, with the most pronounced differences observed during the initial uptake phase (0.5 h) between these two phagocytic pathways (Fig. 4B). We then built phosphosignaling networks from KiR-predicted kinases that were unique to uptake of amastigotes via a C3bi-mediated mechanism, but were not involved in either of the bead phagocytosis pathways. In addition to MAPK signaling pathways, multiple other kinase-kinase connections were implicated in the network built during initial uptake (0.5 h) (Fig. 4C). For instance, network analysis suggested that AKT serine/threonine kinase 1 (AKT1) and glycogen synthase kinase 3 beta (GSK3β) signaling is important for amastigote-initiated infection via a C3bi-mediated mechanism. Interestingly, previous studies have shown that activation of GSK3β suppresses IL-10 production for a protective immune response against *Leishmania* infection [32, 33]. Our model also predicted that SRC-mediated signaling associated with transforming growth factor beta receptor (TGFβ-R) is essential for uptake of amastigotes via binding to CR3. Consistent with this prediction, both field and *in vitro* studies conducted by others have reported that TGFβ-Rs are involved in *Leishmania* infection, although these were done with *Leishmania* species other than *L. amazonensis* [34, 35]. Overall and in summary, our results demonstrate the potential of using KiR and network-level analysis to systematically dissect key kinase regulators and underlying signaling pathways associated with *Leishmania* infection.

**Fig. 4.**
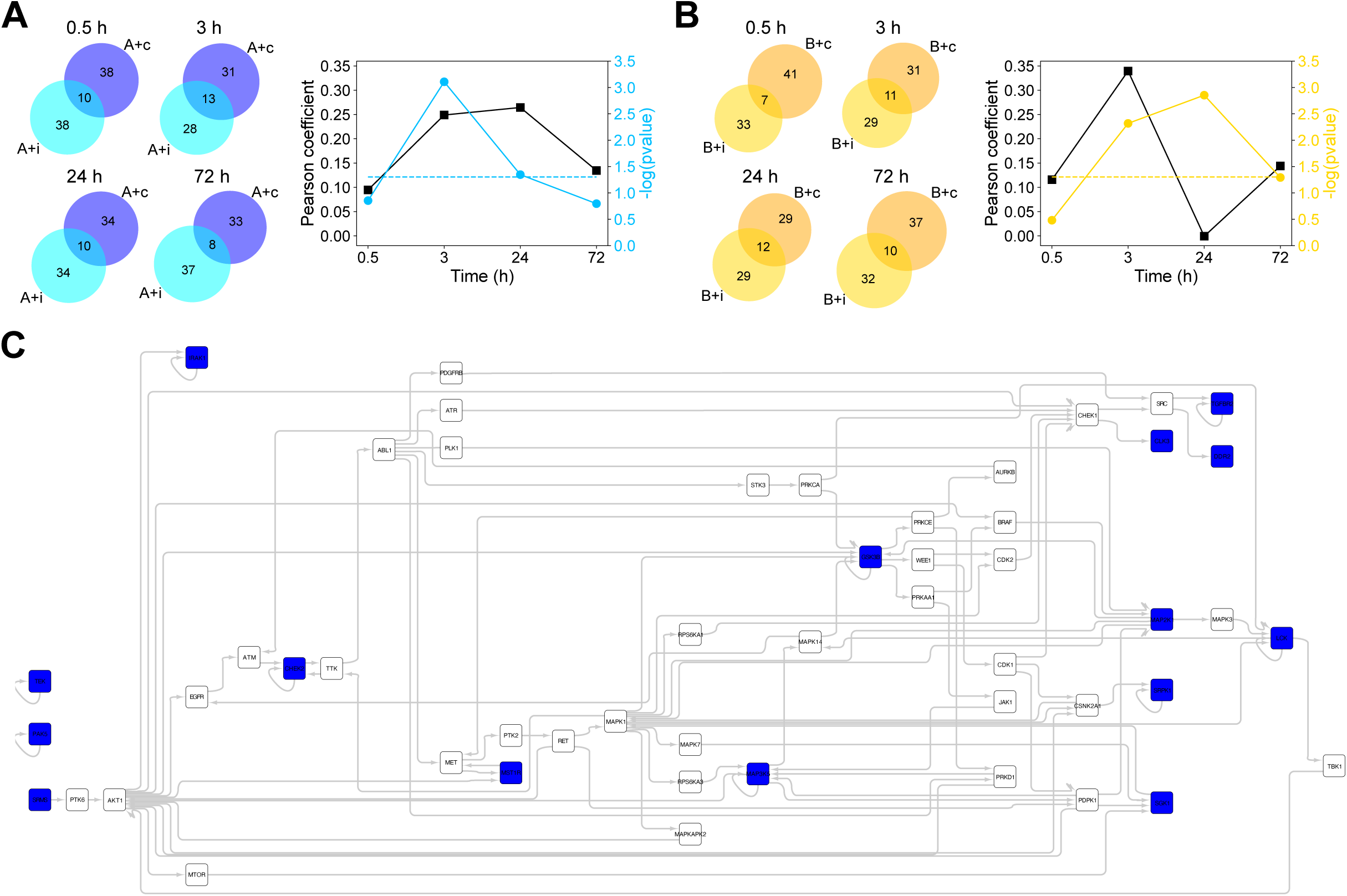
Host kinases differentially regulate *Leishmania* infection via C3bi- and IgG-mediated entry mechanisms. **(A)** Left: Venn diagrams showing the number of kinases predicted to modulate C3bi- and IgG-mediated uptake of amastigotes at different stages of infection. Right: black squares represent the Pearson correlation coefficients calculated from the normalized phagocytic index upon kinase inhibitor treatment between C3bi- and IgG-mediated uptake of amastigotes; light blue circles represent the p-values from hypergeometric test performed on KiR-predicted kinases between C3bi- and IgG-mediated uptake of amastigotes; light blue dashed line corresponds to p-value of 0.05. **(B)** Left: Venn diagrams showing the number of kinases predicted to modulate C3bi- and IgG-mediated uptake of beads at different time points. Right: black squares represent the Pearson correlation coefficients calculated from the normalized phagocytic index upon kinase inhibitor treatment between C3bi- and IgG-mediated uptake of beads; gold circles represent the p-values from hypergeometric test performed on KiR-predicted kinases between C3bi- and IgG-mediated uptake of beads; gold dashed line corresponds to p-value of 0.05. **(C)** Phosphosignaling network built from the kinases predicted to exclusively regulate amastigote-initiated infection via a C3bi-mediated mechanism at 0.5 h post infection (labeled in blue), using the kinase-substrate phosphorylation database on PhosphoSitePlus^®^ as the background network.

## 4. Discussion

Here, we implemented an image-based phenotypic screen and elastic net regression algorithm to propose host kinase regulators of *L. amazonensis* promastigote and amastigote uptake and survival within macrophages. Furthermore, to inform the field regarding which kinases could be required for general phagocytic mechanisms, we performed the same experiments using coated inert beads. Our findings are presented as a tool for the scientific community to engender further studies of *Leishmania* uptake by, and survival within, mammalian macrophages. Our approach identifies several candidate host kinases regulating receptor-mediated phagocytosis and/or the uptake of *Leishmania*. In addition, we delineate putative kinases necessary for promastigotes to convert to amastigotes or for amastigotes to survive within macrophages. Our data suggest that the host kinases required for these processes differ depending on which *Leishmania* life cycle stage is internalized and which cell surface receptors are bound. Our data also indicate that uptake of *Leishmania* parasites by macrophages differ substantially from generalized phagocytic mechanisms.

We initially used 38 inhibitors to delineate kinases that affect *Leishmania* or bead internalization, as well as *Leishmania* survival inside macrophages. Although there were variations between phagocytic indexes and surviving parasites over biological replicates, we were still able to identify several kinases that have been implicated in these processes. For example, both Abl and Src inhibitors prevented *Leishmania* internalization and phagocytosis in our assay, as we have shown previously [13, 14]. Furthermore, we documented kinases affecting C3bi- and IgG-mediated bead uptake that have previously been implicated in phagocytosis, such as ROCK2 [29]. Overall, these results help to internally validate our kinase screen. Importantly, our results suggested many other kinases that could be involved in *Leishmania* uptake, such as MAP kinases and WNK2, which we are testing in our ongoing and published work [17]. Admittedly, we do not wash away inhibitors during our assays, so the 38 kinase inhibitors we tested technically could affect parasites as well as host cells. However, our ongoing and future validation efforts for the kinases we identified will test for any effects of inhibitors on parasites themselves.

We then used a machine learning method to predict additional kinases required for either uptake of parasites or beads, or subsequent survival of *Leishmania* within macrophages. Although the KiR methodology does not consider factors such as protein abundance, subcellular localization, translation, degradation, and dephosphorylation, which are all vital for comprehending cell signaling driven by post-translational modifications, it provides the capability to link functional kinases with cellular phenotypes in an unbiased and systematic manner. By building phosphosignaling networks, we expanded the scope from the 291 kinases in the biochemical dataset used for KiR to the entire human kinome, and connected functional kinases within a network structure that describes the pathways through which phosphosignals propagate. Thus, our approach provides insights on the underlying kinase signaling mechanisms controlling *Leishmania* uptake and intracellular development.

Established literature would suggest that *Leishmania* uptake by macrophages involves generalized phagocytic mechanisms [9], even though this complex organism possesses substantial structural and proteomic differences from other commonly-phagocytosed particles. However, our model predicted that kinases and kinase signaling pathways regulating *Leishmania* uptake by macrophages would be substantially different than those required for phagocytosis of coated beads. Our ongoing studies presented here and elsewhere have begun to experimentally validate this finding. For example, in Fig 2F, we see some differences in the degree to which MAP kinase inhibition limits the uptake of amastigotes compared to beads; see also [17]). Thus, our findings challenge prior hypotheses in the field and suggest that further studies of the internalization process will not only uncover novel mechanisms for *Leishmania* entry into macrophages, but could also suggest novel host-active agents that could be specific to treat leishmaniasis.

Furthermore, our studies suggest that host kinase signaling pathways change over time as *Leishmania* parasites establish their intracellular niche, which has been demonstrated by others [9]. However, and of significant interest, our data suggest that even signaling events that are related to parasite survival and subsequent development taking place hours to days after internalization can be influenced by the initial receptor that is bound. Our results both complement and add to prior literature suggesting that activating different cell surface receptors on phagocytes affects the eventual fate of this intracellular parasite.

In addition, we find that different life cycle stages of *L. amazonensis* may trigger different host kinase signaling events during their uptake by and survival within macrophages. Given the remarkable differences between parasite size and composition in these two life cycle stages, perhaps this is not altogether surprising. However, we simultaneously find that the host macrophage receptor to which a *Leishmania* amastigote initially binds may often affect later signaling events while the parasite resides within the macrophage. We anticipate that in turn, these differential signaling pathways would then affect the parasite’s eventual fate and survival, which we will test directly in our future work.

In conclusion, our studies implicate many candidate host kinases in *L. amazonensis* promastigote and amastigote internalization by macrophages as well as generalized phagocytosis. We have also proposed host kinase signaling pathways seen while promastigotes convert to amastigotes within the macrophage or are activated while *Leishmania* survives within the acidic phagolysosome. Interestingly, it appears there are likely many differences between the kinases that play roles in *Leishmania* versus bead internalization, promastigote versus amastigote uptake and survival, and CR3 versus FcR-activated pathways. As these mechanisms may also differ based on the infecting species of *Leishmania*, further studies will be directed at identifying differences in kinase signaling pathways between entry and survival of these life cycle stages in different parasite species. Uncovering the kinases that control the internalization of *Leishmania* by macrophages and its subsequent growth and development in phagocytes may lead to novel host-based targets for new antimicrobials.

## Data availability

Data generated in this study are available in the main text or the supplementary information.

## Acknowledgements

We would like to thank members and colleagues of the Wetzel and Kaushansky laboratories for technical assistance and helpful comments, including Laela M. Booshehri, James M. Bradford, Lauren T. Callaghan, Ivan Luu, Emily T. Mamula, Emma L. Rhodes, Catherine Trice, and Imran Ullah. Drs. Hanspeter Niederstrasser and Bruce A. Posner in the UT Southwestern High Throughput Screening Core helped image the initial uptake and survival kinase inhibitor screen with the Core’s InCell Analyzer 6000, which was purchased through NIH S10 OD018005. This work has been supported by NIH K08 AI103036, NIH R01 AI146349, Children’s Clinical Research Advisory Committee (CCRAC) Junior and Early Investigator Awards, a Welch Grant for Chemistry (I-2086), and funds from the Department of Pediatrics (all to DMW), and NIH R01 GM101183 (to AK). UB and GMA were supported by Medical Scientist Training (MD/PhD) Grant NIH T32 GM008014. In addition, UB was funded by a National Institutes of Health Supplement to Promote Diversity in Health-Related Research (R01 AI146349-S1).

## CRediT authorship contribution statement

**Ling Wei:** Conceptualization, Investigation, Writing – Original Draft, Writing – Review & Editing. **Umaru Barrie:** Investigation, Validation, Formal Analysis, Data Curation, Visualization, Writing – Original Draft, Writing – Review & Editing. **Gina M. Aloisio:** Methodology, Validation, Investigation, Formal Analysis. **Francis T. H. Khuong:** Validation, Formal Analysis, Data Curation, Investigation, Writing – Review & Editing. **Nadia Arang:** Conceptualization, Formal Analysis. **Arani Datta:** Validation, Formal Analysis, Data Curation, Investigation, Writing – Review & Editing. **Alexis Kaushansky:** Conceptualization, Funding Acquisition, Supervision, Project Administration, Writing – Review & Editing. **Dawn M. Wetzel:** Conceptualization, Methodology, Formal Analysis, Resources, Funding Acquisition, Supervision, Project Administration, Writing – Original Draft, Writing – Review & Editing.

## Declaration of Competing Interest

The authors declare that they have no competing interests.

## Supplemental Figures

**Fig. S1. Correlations of normalized phagocytic index upon kinase inhibitor treatment between different conditions. (A)** Cluster map showing the impact of a panel of 38 kinase inhibitors on *Leishmania* infection of macrophages. Phagocytic indices for kinase inhibitor-treated cells were normalized to the phagocytic index of DMSO-treated control cells. Three independent experiments were performed. **(B)** Correlation between amastigote and bead uptake via an IgG-mediated mechanism. **(C)** Correlation between promastigote and bead uptake via a C3bi-mediated mechanism. **(D)** Correlation between amastigote and bead uptake via a C3bi-mediated mechanism. **(E)** Correlation of bead uptake between IgG- and C3bi-mediated mechanisms. Correlations were calculated using the mean values of all three biological replicates. Pearson correlation coefficient (r) is shown in each plot.

**Fig. S2. Cluster map of the residual activity of 291 protein kinases targeted by the 38 kinase inhibitors in screens.** Residual kinase activity data are from Anastassiadis *et al*. The impact of the kinase inhibitors on *Leishmania* infection in macrophages is shown as a colormap at bottom.

**Fig. S3. Phosphosignaling networks built from the overlapping kinases predicted in C3bi-mediated uptake of promastigotes and beads.** Networks were built at 3 h (A) and 0.5 h (B) post internalization. Kinases predicted by KiR are in orange.

## Supplemental Tables

**Table S1. Kinase-compound biochemical data and cognate targets of the 38 kinase inhibitors used in this screen.** Sheet “biochemical” contains residual kinase activity data of the 38 kinase inhibitors used in this screen against 291 recombinant kinases; this data is from Anastassiadis *et al* [30]. Sheet “cognate” contains the lists of cognate targets of these kinase inhibitors; information is from DrugBank or vendors’ websites. This is a Microsoft Excel workbook containing 2 spreadsheets.

**Table S2. Normalized phagocytic indexes (with respect to DMSO (100%)) from kinase inhibitor screens.** Left: Three biological replicates were performed at each condition. Right: Normalized mean phagocytic index (PI) and standard error for each inhibitor at each time point. This is a Microsoft Excel workbook containing 5 spreadsheets.

**Table S3. List of kinases predicted by KiR.** Kinases predicted to inhibit or promote uptake/survival are in blue or red, respectively. This is a Microsoft Excel workbook containing 5 spreadsheets.

## Notes

### Competing Interest Statement

The authors have declared no competing interest.

### Summary of Updates

We have included additional analysis of our data and revised our phrasing.

## References

1. Burza, S., S.L. Croft, and M. Boelaert, Leishmaniasis. Lancet, 2018. 392(10151): p. 951–970.

2. CDC. Parasites - Leishmaniasis. 2022; Available from: https://www.cdc.gov/parasites/leishmaniasis/index.html.

3. Sibley, L.D. and N.W. Andrews, Cell invasion by un-palatable parasites. Traffic, 2000. 1(2): p. 100–6.

4. Kane, M.M. and D.M. Mosser, Leishmania parasites and their ploys to disrupt macrophage activation. Curr Opin Hematol, 2000. 7(1): p. 26–31.

5. Wenzel, U.A., et al., Leishmania major parasite stage-dependent host cell invasion and immune evasion. FASEB J, 2012. 26(1): p. 29–39.

6. Brittingham, A., et al., Interaction of Leishmania gp63 with cellular receptors for fibronectin. Infect Immun, 1999. 67(9): p. 4477–84.

7. Mosser, D.M. and L.A. Rosenthal, Leishmania-macrophage interactions: multiple receptors, multiple ligands and diverse cellular responses. Semin Cell Biol, 1993. 4(5): p. 315–22.

8. Wilson, M.E. and R.D. Pearson, Roles of CR3 and mannose receptors in the attachment and ingestion of Leishmania donovani by human mononuclear phagocytes. Infect Immun, 1988. 56(2): p. 363–9.

9. Ueno, N. and M.E. Wilson, Receptor-mediated phagocytosis of Leishmania: implications for intracellular survival. Trends Parasitol, 2012. 28(8): p. 335–44.

10. Peters, C., et al., The role of macrophage receptors in adhesion and uptake of Leishmania mexicana amastigotes. J Cell Sci, 1995. 108 (Pt 12): p. 3715–24.

11. Morehead, J., I. Coppens, and N.W. Andrews, Opsonization modulates Rac-1 activation during cell entry by Leishmania amazonensis. Infect Immun, 2002. 70(8): p. 4571–80.

12. Wetzel, D.M., et al., Actin filament polymerization regulates gliding motility by apicomplexan parasites. Mol Biol Cell, 2003. 14(2): p. 396–406.

13. Wetzel, D.M., D. McMahon-Pratt, and A.J. Koleske, The Abl and Arg kinases mediate distinct modes of phagocytosis and are required for maximal Leishmania infection. Mol Cell Biol, 2012. 32(15): p. 3176–86.

14. Datta, A., U. Barrie, and D.M. Wetzel, A Multi-Color Immunofluorescence Assay to Distinguish Intracellular From External Leishmania Parasites. Bio Protoc, 2024. 14(11): p. e5009.

15. Wetzel, D.M., et al., The Src kinases Hck, Fgr, and Lyn activate Abl2/Arg to facilitate IgG-mediated phagocytosis and Leishmania infection. J Cell Sci, 2016. 129(16): p. 3130–43.

16. Ullah, I., et al., Src- and Abl-family kinases activate spleen tyrosine kinase to maximize phagocytosis and Leishmania infection. J Cell Sci, 2023. 136(14).

17. Barrie, U., et al., MAPK/ERK activation in macrophages promotes Leishmania internalization and pathogenesis. Microbes Infect, 2024. 26(5-6): p. 105353.

18. Arang, N., et al., Identifying host regulators and inhibitors of liver stage malaria infection using kinase activity profiles. Nat Commun, 2017. 8(1): p. 1232.

19. Dankwa, S., et al., Exploiting polypharmacology to dissect host kinases and kinase inhibitors that modulate endothelial barrier integrity. Cell Chem Biol, 2021.

20. Wei, L., et al., Interrogating endothelial barrier regulation by temporally resolved kinase network generation. Life Sci Alliance, 2024. 7(5).

21. Bourgeois, N.M., et al., Multiple receptor tyrosine kinases regulate dengue infection of hepatocytes. Front Cell Infect Microbiol, 2024. 14: p. 1264525.

22. Gujral, T.S., L. Peshkin, and M.W. Kirschner, Exploiting polypharmacology for drug target deconvolution. Proc Natl Acad Sci U S A, 2014. 111(13): p. 5048–53.

23. Bates, P.A., et al., Axenic cultivation and characterization of Leishmania mexicana amastigote-like forms. Parasitology, 1992. 105 (Pt 2): p. 193–202.

24. Bates, P.A., Complete developmental cycle of Leishmania mexicana in axenic culture. Parasitology, 1994. 108 (Pt 1): p. 1–9.

25. Russell, D.G. and S.D. Wright, Complement receptor type 3 (CR3) binds to an Arg-Gly-Asp-containing region of the major surface glycoprotein, gp63, of Leishmania promastigotes. J Exp Med, 1988. 168(1): p. 279–92.

26. Guy, R.A. and M. Belosevic, Comparison of receptors required for entry of Leishmania major amastigotes into macrophages. Infect Immun, 1993. 61(4): p. 1553–8.

27. Dixit, U.G., et al., Complement receptor 3 mediates ruffle-like, actin-rich aggregates during phagocytosis of Leishmania infantum metacyclics. Exp Parasitol, 2021. 220: p. 107968.

28. Greuber, E.K. and A.M. Pendergast, Abl family kinases regulate FcgammaR-mediated phagocytosis in murine macrophages. J Immunol, 2012. 189(11): p. 5382–92.

29. Barger, S.R., N.C. Gauthier, and M. Krendel, Squeezing in a Meal: Myosin Functions in Phagocytosis. Trends Cell Biol, 2020. 30(2): p. 157–167.

30. Anastassiadis, T., et al., Comprehensive assay of kinase catalytic activity reveals features of kinase inhibitor selectivity. Nat Biotechnol, 2011. 29(11): p. 1039–45.

31. Hornbeck, P.V., et al., PhosphoSitePlus, 2014: mutations, PTMs and recalibrations. Nucleic Acids Res, 2015. 43(Database issue): p. D512–20.

32. Paul, J., et al., TLR mediated GSK3beta activation suppresses CREB mediated IL-10 production to induce a protective immune response against murine visceral leishmaniasis. Biochimie, 2014. 107 Pt B: p. 235-46.

33. Nandan, D., et al., Myeloid cell IL-10 production in response to leishmania involves inactivation of glycogen synthase kinase-3beta downstream of phosphatidylinositol-3 kinase. J Immunol, 2012. 188(1): p. 367–78.

34. Weirather, J.L., et al., Comprehensive candidate gene analysis for symptomatic or asymptomatic outcomes of Leishmania infantum infection in Brazil. Ann Hum Genet, 2017. 81(1): p. 41–48.

35. Srivastava, S., et al., Leishmania expressed lipophosphoglycan interacts with Toll-like receptor (TLR)-2 to decrease TLR-9 expression and reduce anti-leishmanial responses. Clin Exp Immunol, 2013. 172(3): p. 403–9.

