## Supplementary figures and images for "Using machine learning to dissect host kinases required for *Leishmania* internalization and development"

### Supplemental Figure 1

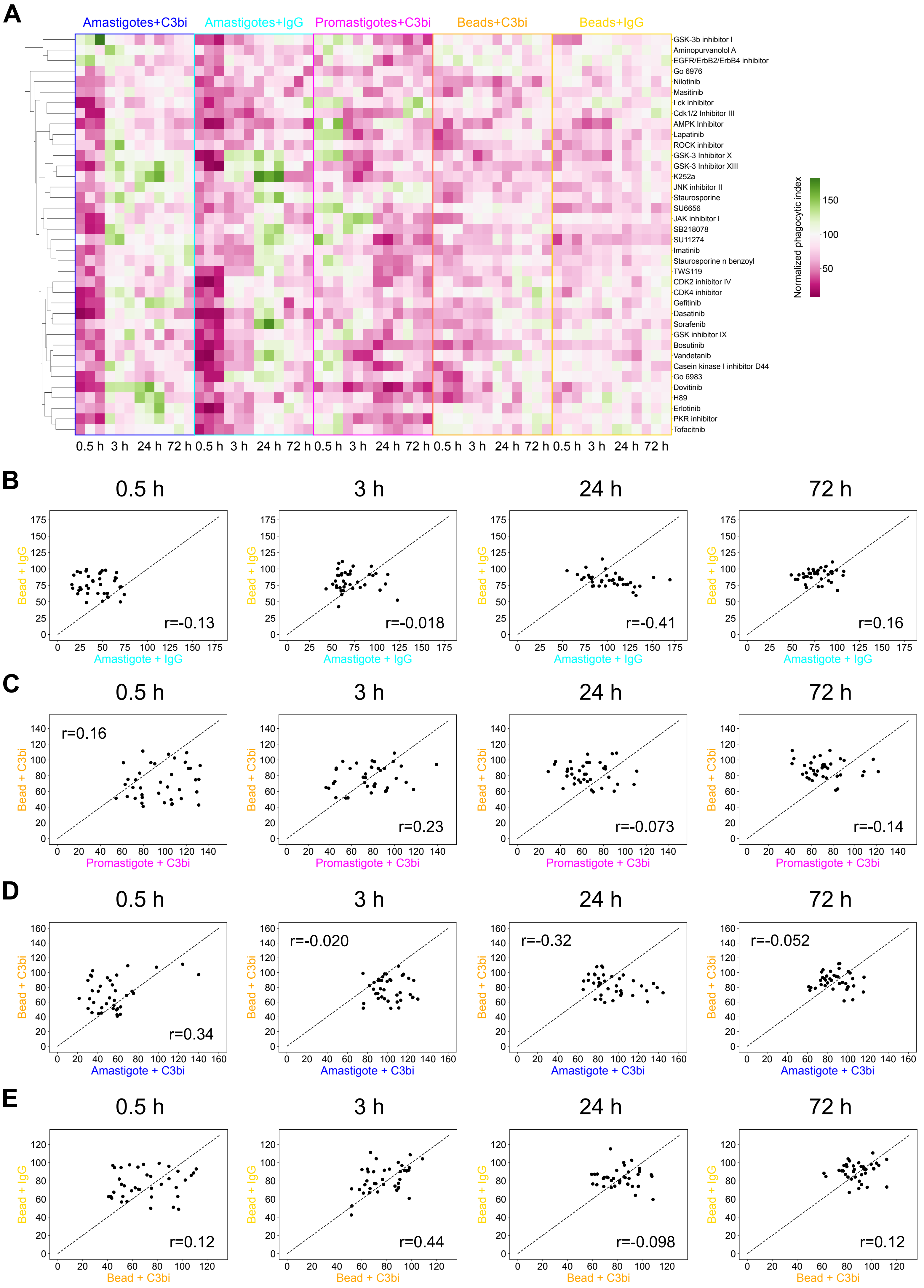

### Supplemental Figure 2

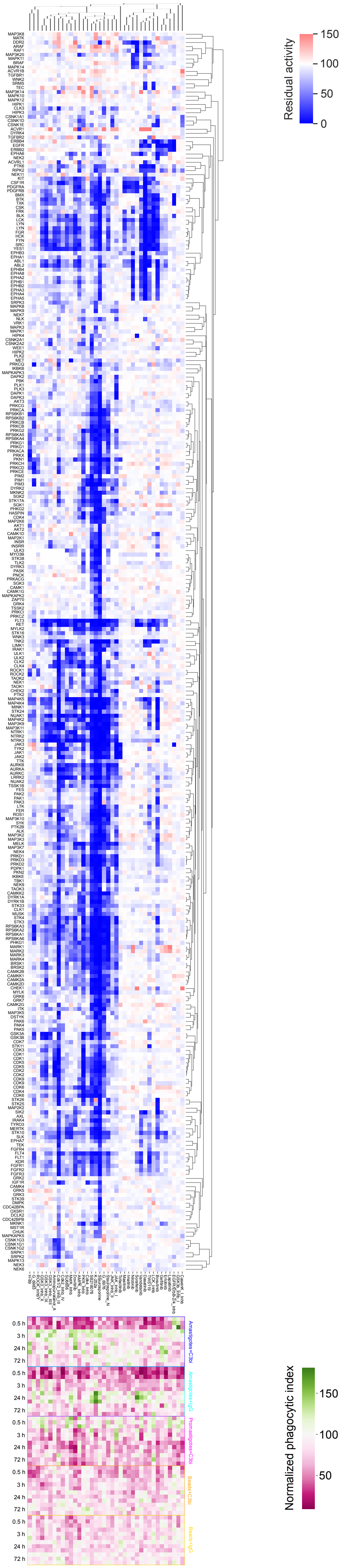

### Supplemental Figure 3

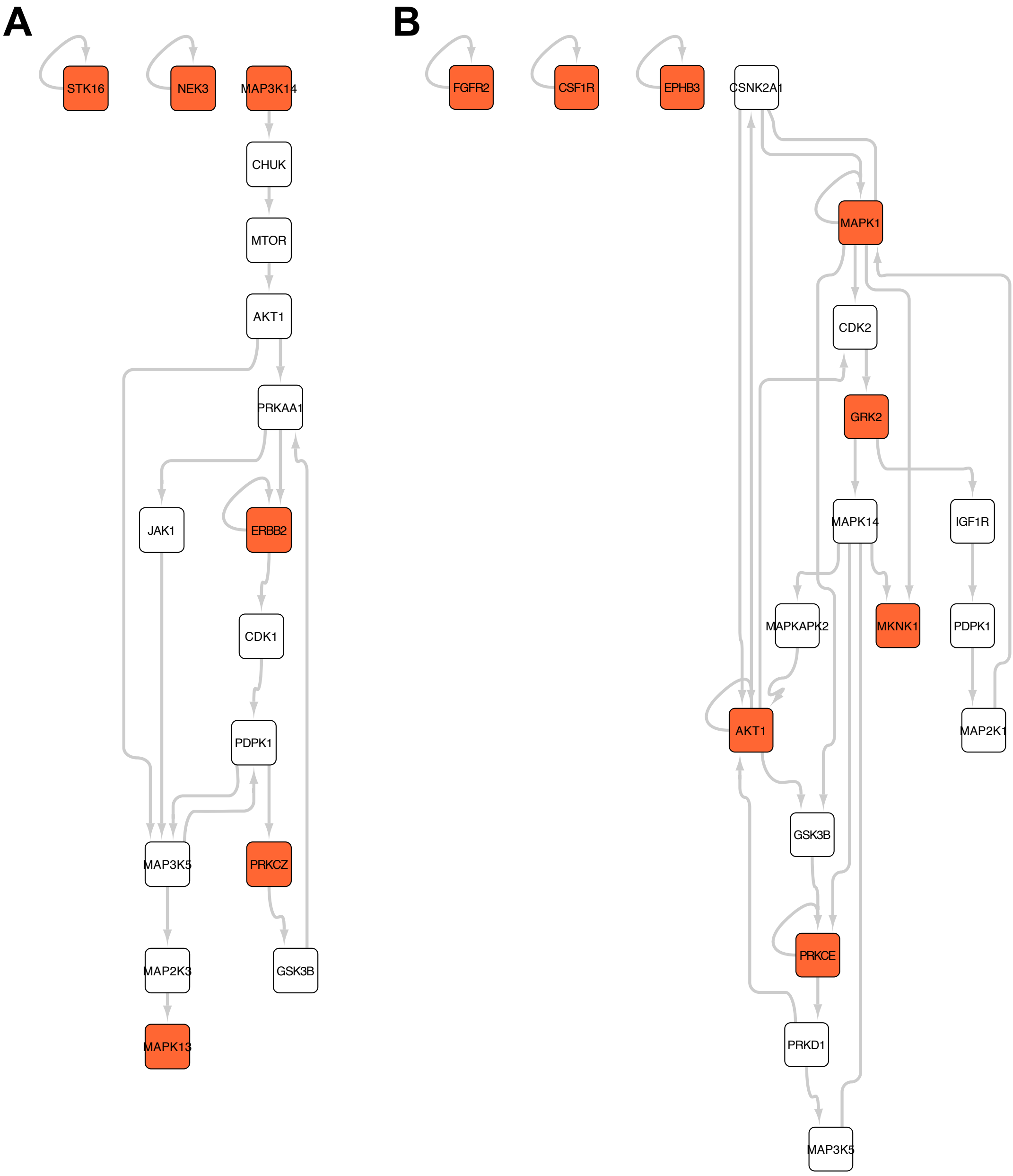
